# Recapitulation of patient-specific 3D chromatin conformation using machine learning and validation of identified enhancer-gene targets

**DOI:** 10.1101/2021.11.16.468857

**Authors:** Duo Xu, Andre Neil Forbes, Sandra Cohen, Ann Palladino, Tatiana Karadimitriou, Ekta Khurana

**Affiliations:** Sandra and Edward Meyer Cancer Center, Weill Cornell Medicine, New York, NY, USA; Institute for Computational Biomedicine, Weill Cornell Medical College, New York, New York, USA; Department of Physiology and Biophysics, Weill Cornell Medical College, New York, New York, USA; Englander Institute for Precision Medicine, Weill Cornell Medicine, New York, NY, USA; Weill Cornell Graduate School of Medical Sciences, Weill Cornell Medicine, New York, New York, USA

## Abstract

Regulatory networks containing enhancer to gene edges define cellular state and their rewiring is a hallmark of cancer. While efforts, such as ENCODE, have revealed these networks for reference tissues and cell-lines by integrating multi-omics data, the same methods cannot be applied for large patient cohorts due to the constraints on generating ChIP-seq and three-dimensional data from limited material in patient biopsies. We trained a supervised machine learning model using genomic 3D signatures of physical enhancer-gene connections that can predict accurate connections using data from ATAC-seq and RNA-seq assays only, which can be easily generated from patient biopsies. Our method overcomes the major limitations of correlation-based approaches that cannot distinguish between distinct target genes of given enhancers in different samples, which is a hallmark of network rewiring in cancer. Our model achieved an AUROC (area under receiver operating characteristic curve) of 0.91 and, importantly, can distinguish between active regulatory elements with connections to target genes and poised elements with no connections to target genes. Our predicted regulatory elements are validated by multi-omics data, including histone modification marks from ENCODE, with an average specificity of 0.92. Application of our model on chromatin accessibility and transcriptomic data from 400 cancer patients across 22 cancer types revealed novel cancer-type and subtype-specific enhancer-gene connections for known cancer genes. In one example, we identified two enhancers that regulate the expression of ESR1 in only ER+ breast cancer (BRCA) samples but not in ER-samples. These enhancers are predicted to contribute to the high expression of ESR1 in 93% of ER+ BRCA samples. Functional validation using CRISPRi confirms that inhibition of these enhancers decreases the expression of ESR1 in ER+ samples.

## Introduction

Enhancers are non-coding regions that play an important role in gene regulation. Transcriptional reprogramming accompanied by enhancer rewiring is a hallmark of cancer. Most studies to characterize enhancers in cancer have used functional genomics data from cell lines or mouse models. This is because enhancer identification generally requires ChIP-seq assays, or transcription-based approach (Cap analysis of gene expression, CAGE), which need larger amounts of input material (∼10^7^ cells) than available from patient biopsies (∼10^6^ cells or less) (*1-5*). However, the advent of Assay for Transposase Accessible Chromatin using Sequencing (ATAC-Seq), which can be performed with lower input material, has led to the identification of open chromatin regions in individual patient samples (*1, 6, 7*). Chromatin accessibility is a key feature of actively transcribed regions and regulatory elements because open chromatin is a prerequisite for transcription factors (TFs) and chromatin regulators (CRs) to bind (*8, 9*). It is known that not all the open chromatin regions are enriched by functional marks for regulatory elements (*2*). Moreover, open chromatin regions may correspond to different regulatory elements, such as, enhancers, promoters, silencers or insulators (*5, 10, 11*). Furthermore, these elements may also be in different states, e.g. for enhancers, active vs. poised (*11, 12*). Such complexity and dynamic regulatory context further complicate the identification of active regulatory elements solely based on ATAC-seq.

Besides the identification of enhancer regions, identification of their target genes has remained a challenge. In the past five years, multiple methods have been used to connect regulatory elements to their target genes both experimentally and computationally for representative tissues and cell lines. Experimentally, cohesin ChIA-PET and H3K27ac HiChIP have been applied to various cell types to discover physical contacts of regulatory elements to target genes (*13, 14*). CRISPR interference (CRISPRi) screening has also been applied to individual genes to identify enhancer-gene connections (*15*). Moreover, hybrid methods integrating CRISPRi-FlowFISH with ChIP-seq and Hi-C (*16*) or comprehensive sets of histone modification ChIP-seq with machine learning (*17*) have also been used to identify enhancer-gene connections for representative tissues and cell lines. However, these experiments are expensive and difficult to perform at scale, especially given the aforementioned limits on input sample. While previous efforts have used correlations between ATAC-seq signal and nearest gene expression across samples to identify target genes (*18*), it has been shown that the 1) the regulatory elements don’t always regulate the closest gene (*2*); 2) the regulatory relationship between regulatory element and gene varies among different cells or tissue types and is also expected to vary among patient samples. Thus, global correlations of element-gene connections may not be able to identify cell-type-specific regulation or combinatorial effects of multiple regulatory elements on expression (*14, 19*); 3) when assigning element-gene connections at patient-specific level, correlation based methods are limited by assigning the connection based on the chromatin opening or not. However, it has been known that not all open regions are active regulatory elements promoting gene expression, there are open chromatin sites that can be bound by repressive protein complexes to repress the gene expression (*11, 12*). Thus, the correlation methods would detect more false positive connections on a per patient basis.

To resolve the above issues, we developed DGTAC (Differential Gene Targets of Accessible Chromatin), which can identify target genes of accessible chromatin regions in a sample-specific manner. DGTAC is trained using 3D conformation data to predict enhancer to gene links using only ATAC-seq and RNA-seq data. The power of sample-specific prediction is demonstrated by a) its ability to distinguish active vs. poised enhancer states, and b) identification of novel enhancers that act in a subtype specific manner to regulate key cancer genes. Application of DGTAC on chromatin accessibility and transcriptomic data from 371 cancer patients of 22 cancer types revealed novel cancer-type and -subtype-specific enhancer-gene connections for 637 known cancer genes. Our predictions can be validated by existing ChIP-seq datasets and experiment-based (HiChIP, Hi-C) or hybrid methods (Activity-by-contact method) predicted connections. An important example is the identification of two enhancers that may explain the upregulation of ESR1 in 93% of ER-positive (ER+) breast cancer samples and are either not accessible or exist in a poised state in ER-negative (ER-) samples. Functional validation using CRISPRi confirms that inhibition of these enhancers decreases the expression of ESR1. Thus, DGTAC can identify the complex architecture of gene regulation by enhancers in cancer patient samples and it is available as a tool that can be used on any sample with matching ATAC-Seq and RNA-Seq.

## Results

### DGTAC: A machine learning model trained using 3D conformation data to identify patient-specific functional regulatory interactions

Previous studies have shown that regulatory elements and their target TSS need to reside in the same loops insulated by the cohesin/CTCF protein complexes to interact with each other (20, 21). To define the gold standard for the actual physical contacts between peak-gene, we used the cohesin/CTCF ChIA-PET loops. Peak signal has been associated with gene expression in a one-to-one basis to connect the accessible peaks to target genes (1). In contrast to one-to-one correlation, we used ElasticNet regression to correlate all ATAC-seq peaks within the 0.5 Mbp range of each transcription start site (TSS) to its expression. The coefficients of each peak to the expression of TSS are calculated (Figure 1b).

**Figure 1.**
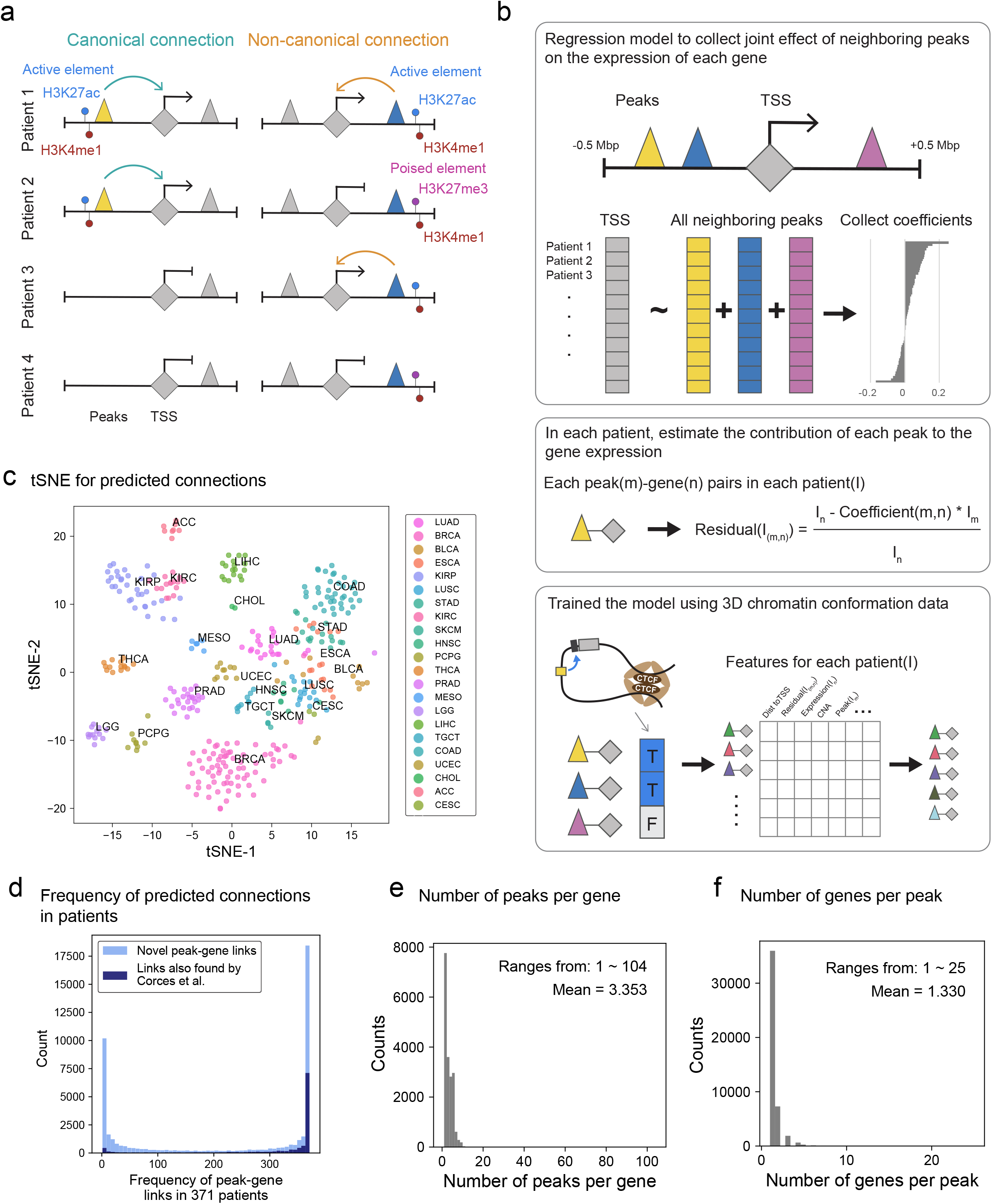
A two-step model to identify patient-specific functional regulatory interactions. a) a schematic of regulatory elements to gene connection predicted by the model in different patients. Triangles stand for ATAC-seq peaks; diamonds stand for TSS of a gene; colored peaks stand for peaks with predicted target genes, and they have different behavior among patients. b) detailed steps for DGTAC. c) clustering of predicted peak-gene connections in each patient by tSNE, each sample is colored by their annotated cancer type. d) frequency of predicted 61,502 peak-gene connections in 371 patients. The darker blue stands for connections also predicted by Corces et al.; the light blue stands for the novel connections predicted by our model. e) histogram of the number of peaks that a gene can be regulated by. f) histogram of the number of target genes that can be regulated by each peak.

Among all potential 6 million peak-gene pairs, 910,116 peak-gene pairs had non-zero coefficients and were used for the next step. Following the calculation of coefficients across samples, we calculated a residual (e) term as the deviation of the peak’s observed contribution to the gene expression from the expected expression in a given sample (Figure 1b). The smaller the residual term, the smaller the deviation from the expected expression for a given peak and link to that gene and vice versa. While the coefficients from the ElasticNet capture the contribution of the peak to the gene expression across samples, the residual terms capture the sample-specific differences. Next, we collected additional information for each peak-gene pair, including the residual terms, distance to TSS, ATAC-seq signal strengths, genomic copy number, and the expression of the target gene to construct a feature matrix for each patient sample. We used the ChIA-PET datasets from MCF-7 (ER+ breast cancer) and HeLa (cervical cancer) (22). We trained two random forest models on patient samples matching the same cancer subtype (LumA/ER+ breast invasive carcinoma, BRCA for MCF-7 and cervical squamous cell carcinoma and endocervical adenocarcinoma, CESC for HeLa) (**Methods**). Both BRCA-model and CESC-model achieved an AUROC of 91.1%. The F1 scores for applying one model to another are 0.927 for BRCA-model and 0.938 for CESC model (Supplementary table 1). This demonstrates that DGTAC can learn the features that capture the functional association between accessible chromatin and gene expression and can be generalized across cancer types.

In total, we identified 61,502 peak-gene connections, which are supported by the expected cancer-type and -subtype classification (Figure 1c, Supplementary figure 1a, 1b and 1c).

Upon comparison of the frequency of novel element-gene connections detected by DGTAC to the ones also detected by simple correlation-based approach (1), we observe that the novel connections are more sample-specific as expected (Figure 1d). Further inspection of features that contribute to the random forest model shows that the “residual” is the most important sample-specific features. We also compared our prediction for basal breast cancer subtypes and connections predicted by the correlation-based method, with an H3K27ac HiChIP dataset for MDA-MB-231 cell line (13). The precision of our prediction is significantly higher compared to the correlation-based method. When taking the union of element-gene connections, DGTAC predicted for all 371 patients, on average, there are 3.35 peaks (min=1, max=104) that regulate a gene, and a peak regulates 1.33 genes (min=1, max=25) (Figure 1e and 1f). Comparing to 5.53 peaks per gene (min=1, max=117) and 1.60 genes per peak (min=1, max=39) by correlation method (1).

### DGTAC identifies non-canonical element to gene connections which differentiate active vs. poised enhancers

Open chromatin has been noted as the “gatekeeper” of regulatory elements in the genome (23), and it can be associated with multiple regulatory elements, for example, enhancer/silencer, promoter, insulator (24, 25), within which they can also be in different states, e.g. for enhancers, active vs. poised (11, 12, 26). This means that not all open chromatin regions are active regulatory elements, and not all of them are active enhancers. Since DGTAC learns the 3D chromatin conformation from CTCF/cohesin insulator loops, we hypothesize that the peak-gene connections we identify are not insulators but active enhancers and promoters.

In addition, we observed that open regions are not always predicted to be linked to target genes. Specifically, the cases where the chromatin is accessible in multiple samples, but only predicted to function as active enhancers with gene targets in a subset of those samples (Figure 1a). For better identification, we specify those cases as “non-canonical” peak-gene connections, and “canonical” connections are the ones where the presence of the peak is sufficient to indicate that a peak-gene connection exists (**Methods**). We further hypothesize that for those non-canonical connections, the open peaks without target gene predicted are in a poised state and are repressed from regulating gene expression (26) (Figure 1a).

#### Validation using data from ENCODE

To test our hypothesis that open chromatin regions with target gene predicted by DGTAC are active regulatory elements in a sample specific manner, we applied DGTAC for cell lines for which ATAC-seq, RNA-seq (in total 11 cell lines) is available (from ENCODE (2), inhouse and other studies (27), details for dataset are listed in Supplementary table 2), and validated the cell line peak-gene prediction from DGTAC for active enhancers using the H3K27ac ChIP-seq of the same cell line from ENCODE (2) (**Methods**). For better differentiation of enhancers, we separated the promoter peak defined as within - 2500/+100bp of transcription start sites (TSS). We observe that for all cell lines tested, among open peaks for each cell line, our predicted active enhancers (open peaks with target gene predicted) show significantly higher H3K27ac enrichment compared to open peaks without target gene predicted (Figure 2a). Among the 11 cell lines, 6 of them have the annotated candidate regulatory elements from SCREEN by ENCODE III where they annotated candidate regulatory elements by using a combination of H3K27ac, H3K4me3, CTCF ChIP-seq and DNase-seq (2), and 3 of them have only the H3K27ac ChIP-seq annotation. We then take active enhancers predicted for those cell lines and compare them to enhancer-like signatures (ELS) annotated by SCREEN. It shows that the predicted active enhancers from our model are enriched for ELS among all open regions (Fisher’s exact P-values <= 8.34e-159, and average odds ratio = 2.30, Figure 2b, **Methods**). We also inspected the enrichment for promoter-like signatures (PLS), similar enrichment is observed too (Figure 2c). We also evaluate the prediction from patient samples by taking union of enhancer-gene connections for cancer types and perform the same comparison to the closest cell line matching from ENCODE. Same enrichments are observed for both H3K27ac ChIP-seq and SCREEN annotated ELS and PLS (Supplementary figure 2a and 2b, **Methods**).

**Figure 2.**
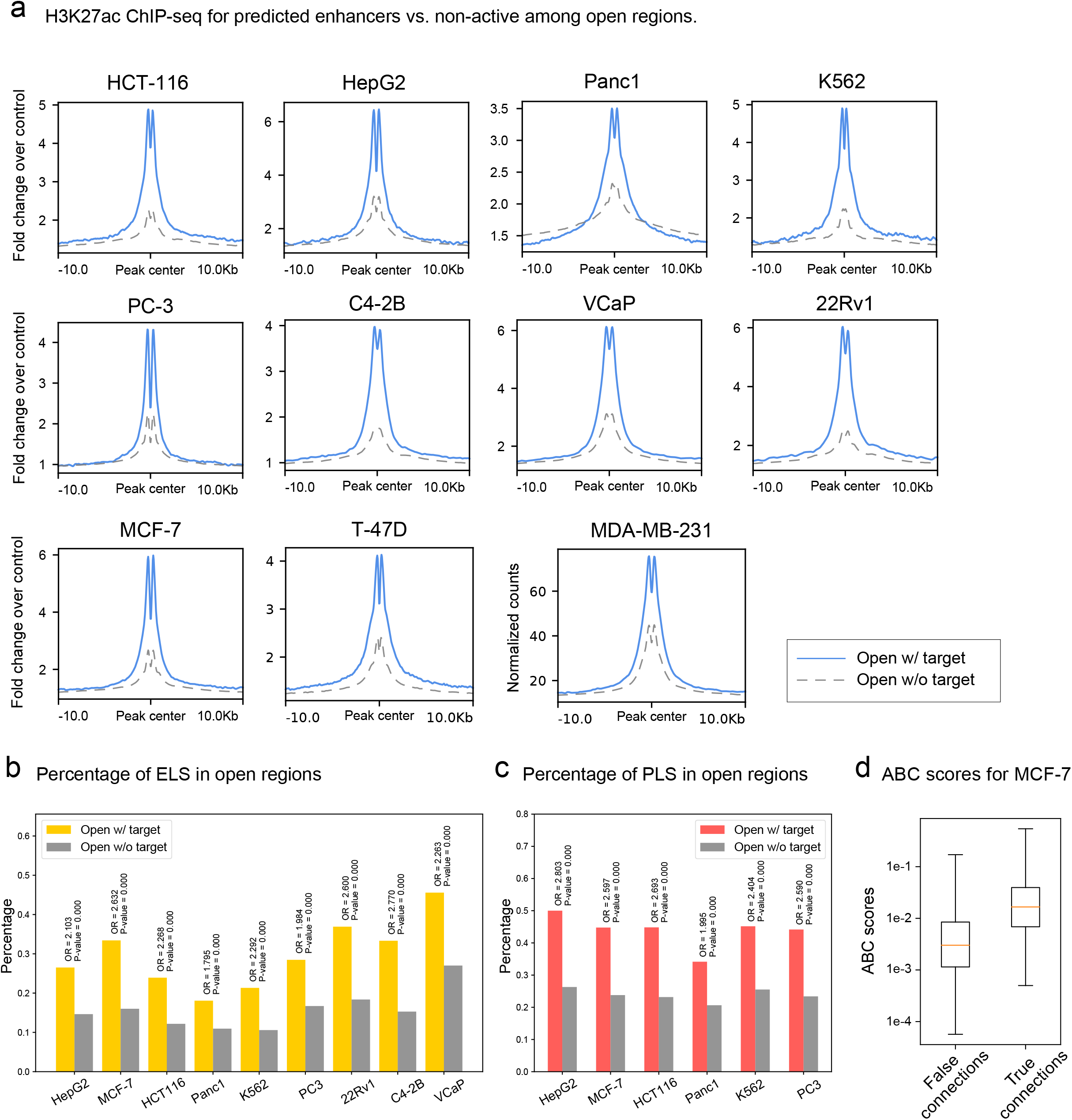
Peaks with target genes predicted are active enhancers. a) For each cell line, the enrichment of H3K27ac ChIP-seq signals comparing between open peaks with target gene predicted vs. open peaks without target gene predicted. b) For cell lines with enhancer or H3K27ac annotation from the SCREEN, among the open peaks, the percentage of enhancer-like signatures (ELS, including both distal-ELS and proximal-ELS) comparing between with vs. without target gene predicted. c) For cell lines with promoter annotation from the SCREEN, among the open peaks, the percentage of promoter-like signatures (PLS) comparing between with vs. without target gene predicted. d) ABC scores of enhancer-gene connections from MCF-7 calculated by us comparing between our predicted True and False connections.

#### Validation using ABC model (MCF-7)

To further validate our prediction of active enhancers, we then compared our prediction to a recently published activity-by-contact model which identifies active enhancers by correlating epigenetic signals (H3K27ac ChIP-seq, DNase-seq, Hi-C) to gene expression perturbed by CRISPRi-FlowFISH (16). Their method uses epigenetic signals to generate an ABC score to measure the element activity, and then correlate the score to expression change by silencing the element to define an ABC score threshold for active enhancers of the assayed cell type. To calculate such an ABC score, we performed ATAC-seq, collected Hi-C from GSE130916 (28) and H3K27ac ChIP-seq from GSM1693017 (29) for MCF-7, and calculated ABC scores for MCF-7 (**Methods**). The calculated ABC scores are significantly higher for connections predicted by DGTAC than ones that are not (Figure 2d, Supplementary figure 2c). If the same cut-off was applied for MCF-7 as provided by Fulco et al. for K562 (cut-off = 0.02) (16), the predicted connections are significantly enriched in connections passing the cut-off (odds ratio = 7.99, Fisher exact P-value < 2.22e-308). These results indicate that without using ChIP-seq and 3D chromatin conformation assays, DGTAC can recapitulate enhancer-gene connections comparable to methods that use a combination of epigenetic assays.

As described above, we observed non-canonical connections in patient samples, and hypothesized that they are “poised” enhancers. Poised enhancers are the ones that after the chromatin becomes accessible, instead of being bound by RNA polymerase II (RNAPII) with tri-acetylation of the H3K27 to initiate the transcriptional activity of the target promoter (30-33), repressive complexes such as polycomb repression complex (PRC2) are recruited, accompanied by tri-methylation of the H3K27, to repress the activity of the target promoter (34). In addition, enhancers can transition between the active and poised states via deacetylation or demethylation, and such transitions are involved in important differentiation processes and tumor stemness. It has been shown in both mice (11) and human embryonic stem cells (10) that poised enhancers can be activated during differentiation (35). This transition was also observed in transcription growth factor β (TGF-β) signaling-triggered epithelial-to-mesenchymal transition (EMT) process which is associated with tumor stemness and metastasis (36).

To test our hypothesis that among the non-canonical connections, the open peaks without target genes are “poised” enhancers, we collected additional ChIP-seq data for histone modification markers such as H3K27me3 for 8 cell lines and compared their enrichment in poised enhancers (open peaks without target genes involving in non-canonical connections) to the active enhancers (open peaks with target genes, Figure 1a). We observed significantly higher enrichment of H3K27me3 in poised vs. active enhancers (Kolmogorov-Smirnov test P-value < 1.35e-18, Figure 3a, Supplementary figure 3a). This observation implies that those open peaks are likely poised enhancers. Further investigation of known pioneering factor FOXA1 and transcription coactivation complex protein P300 from T-47D shows no difference between open peak with and without target genes, further supporting the notion that open peaks without predicted target genes are primed and later poised or silenced (Supplementary figure 3e).

**Figure 3.**
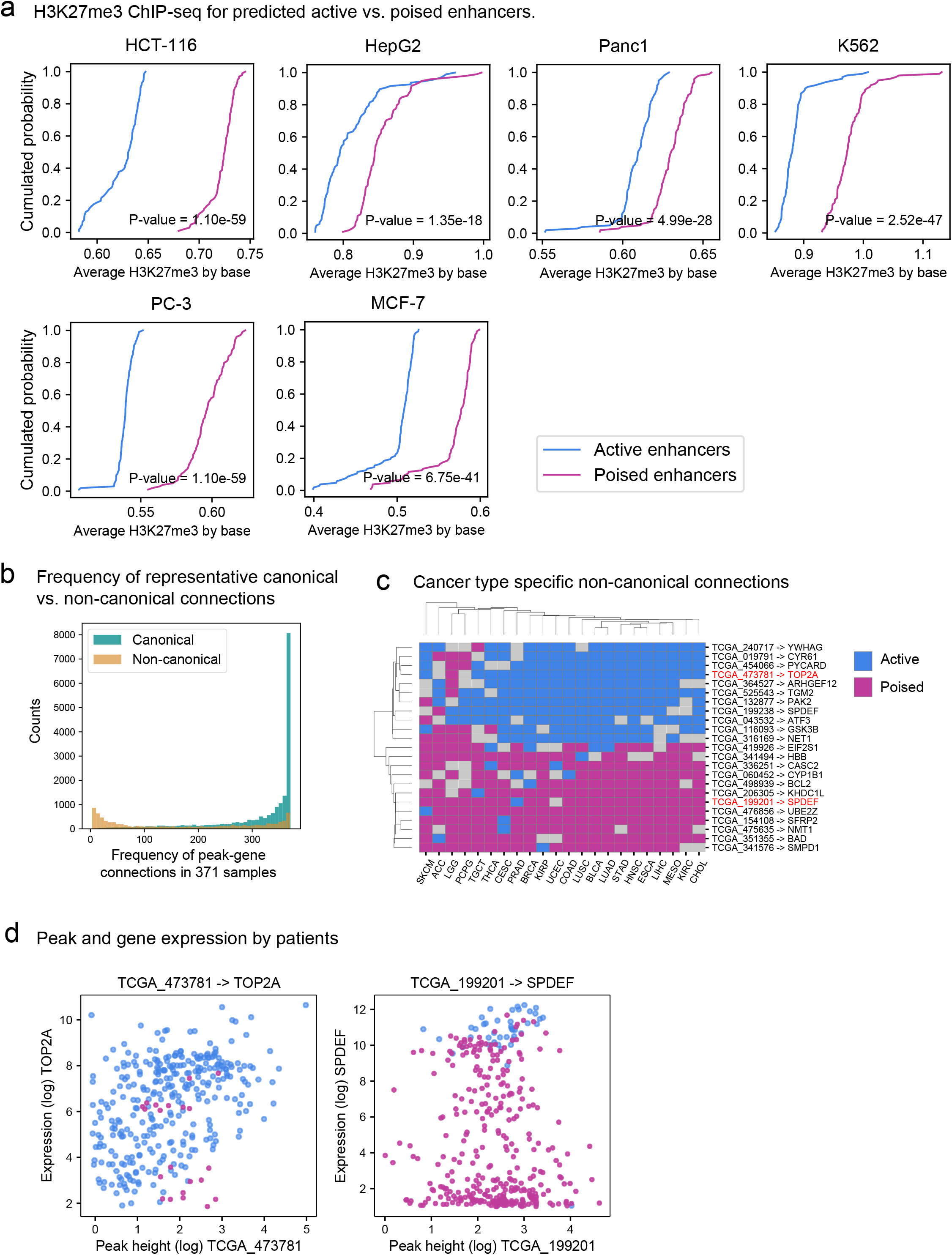
Non-canonical connections reveal poised enhancers. a) Cumulated probability of H3K27me3 ChIP-seq comparing between predicted active vs. poised enhancers. A schema showing canonical vs. non-canonical connections, active vs. poised enhancers and the enriched histone modification makers b) H3K4me1/H3k27ac/H3K27me3 ChIP-seq enrichment for open peaks with vs. without target gene predicted for T-47D and MDA-MB-231. c) FOXA1 and P300 ChIP-seq enrichment for open peaks with vs. without target gene predicted for T-47D. d) Frequency of representative canonical and non-canonical peak-gene connections in 371 patient samples. e) Clustering of representative non-canonical peak-gene connections among patients by tSNE, colored by primary tumor sites. f) Selected cancer type-specifically active vs. poised non-canonical peak-gene connections. Dark yellow vs. purple represent the connection is active or poised in the cancer type respectively. X-axis is cancer types, y-axis is peak-gene connections formatted in “gene-peak”. g) Among the samples with the peak, ESCA_105395, open, the scatter plot for the peak signal (log) and expression of gene TOP2A. Samples with ESCA_105395 L TOP2A connection predicted as active are colored in red, while samples with the connection predicted poised are colored in grey. h) The scatter plot for peak signal of SKCM_38639 and gene expression of SPDEF. Samples with connection predicted as active are colored in red.

Overall, unlike other computational or experimental methods which depend on multiple histone modification makers, especially H3K27ac, to predict active enhancer-gene connections, our model learned the signature of active enhancers from chromatin accessibility and successfully predicted the active enhancers for patients on a large scale using only the ATAC-seq and RNA-seq data.

### Non-canonical connections are more sample-specific and involved in tumorigenesis

To further investigate the non-canonical connections, we curated a group of representative non-canonical peak-gene connections (9,228 connections) with non-canonical score > 0.5. In addition, we also limited the minimum number of samples with or without predicted connections to be more than 5, and the target peak signal should not be significantly different between samples with and without the connection predicted (**Methods**, Supplementary figure 4). In comparison to the non-canonical peak-gene connections, we also curated a group of representative “canonical” peak-gene connections (19,756 connections, **Methods**, Supplementary figure 4, Supplementary table 3). By comparing the frequency of the non-canonical peak-gene connections to canonical ones, we find that non-canonical connections are show lower frequency among 371 samples while most canonical connections are common across samples (Figure 3b). We performed gene ontology enrichment analysis (37) for genes involved in canonical and non-canonical connections comparing them to the genome background. Genes involved in non-canonical connections are enriched in positive regulation of apoptosis or processes associated with apoptosis such as proteolysis and hydrolysis (adjusted P-value < 0.0466, Supplementary table 4), while genes involved in canonical connections are enriched in multiple metabolic and catabolic process (all with adjusted P-value < 7.75e-7). This validates that the enhancer transitions between active vs. poised, which was observed previously in differentiation processes (10, 11, 35) and tumor stemness (36), correspond to non-canonical connections.

**Figure 4.**
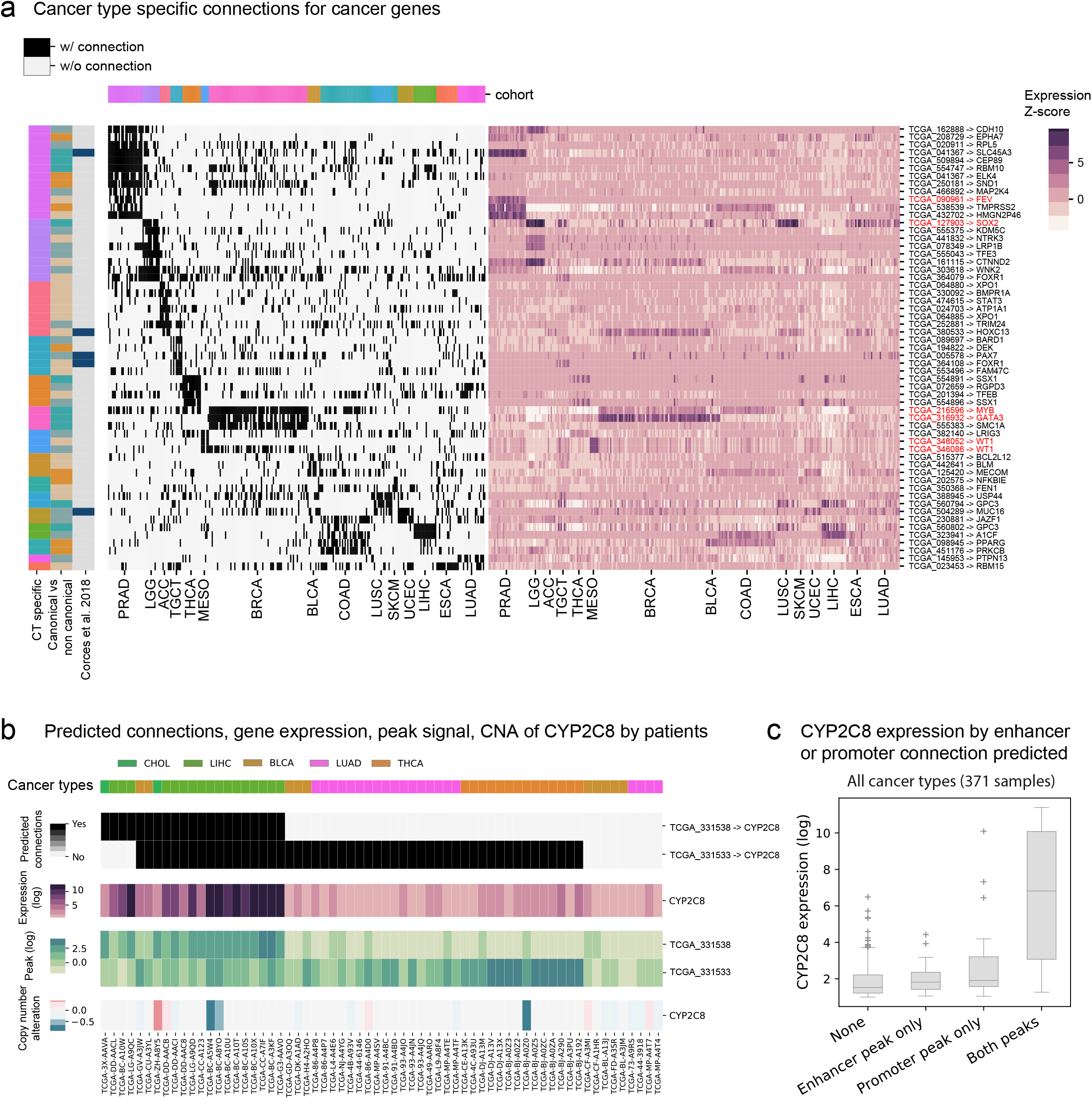
Identification and validation of patient-specific regulatory element to gene interactions. a) cancer-type specific connections by cancer type for cancer genes. The left panel is the connection for each patient sample by cancer types; the right panel is the corresponding expression of genes (log, normalized by medium of the row) involved in the connection for each patient by cancer types. The vertical color bar stands for cancer type specificity of the connection, canonical vs. non-canonical and if the connection was identified from Corces et al. 2018. The horizontal color bar annotates the subtype information for each patient. b) peak-gene connections predicted for CYP2C8 in LIHC/CHOL/BLCA/LUAD/THCA by patients. in addition, expression of CYP2C8 and peak signal for TCGA_331538 (promoter peak) and TCGA_331533 (enhancer peak), as well as copy number alteration for CYP2C8. X-axis is patient sample id. b) boxplot for expression of CYP2C8 from each sample by presence of the enhancer peak (TCGA_331533) CYP2C8 only, promoter peak (TCGA_331538) CYP2C8 only and both peak-gene connections. c) peaks with connections to ESR1 in all samples. Red solid curves are enhancer connections predicted from our model among all sample; dashes grey curves are connections predicted by other studies or random control selected for CRISPRi validation, ATAC-seq signal was from the ATAC-seq sequenced in house for enhDistal and enhIntron respectively.

We evaluated the non-canonical connections involving these apoptosis genes, the frequency of those connections show heterogeneity across cancer types with some connections also showing cancer type specificity that enhancers involved in those connections shows either active or poised state by cancer type (Figure 3c, Supplementary figure 3f, **Methods**). These observations further support the notion that poised enhancers are involved in more cell-type specific regulatory activity, and the tumorigenesis role of enhancers in active or poised state varies across cancer types. For example, one predicted enhancer regulating the TOP2A gene is active in most other cancer types but is poised in brain lower grade glioma (LGG) (Figure 3c and 3d). TOP2A is a known prognostic biomarker for patients with glioma, and its expression is highly positively correlated with glioma grade stage (38). The poised enhancer regulating TOP2A is consistent with the expectation that TOP2A should have low expression in lower grade glioma. As the tumor develops into higher grade glioma, the poised enhancer is acetylated and becomes active, and results in increased expression of TOP2A. Another example is the non-canonical connection for SPDEF, with the enhancer specifically active in prostate adenocarcinoma (PRAD) (Figure 3c and 3d) and correlated with high expression of SPDEF. SPDEF has been known as a tumor suppressor for prostate cancer metastasis (39, 40). It is likely that once the active enhancer becomes poised, as in most other cancers, PRAD patients will lose the expression of SPDEF leading to EMT and bone metastasis (41).

Overall, our model identified a group of non-canonical enhancer-gene connections that are not predicted to target genes despite displaying open chromatin and enhancers without predictions from the non-canonical connections are poised enhancers. Genes involved in non-canonical connections are enriched in apoptosis and the transition from active to poised or from poised to active are heterogenous across cancer types and potentially facilitate tumor metastasis.

### DGTAC identifies novel enhancer-gene connections for cancer genes

Misregulation of key genes is a widely observed hallmark of cancer, thus we wanted to identify enhancer-gene connections that are overrepresented in individual cancers (42-46). To identify cancer-specific peak-gene connections, we first grouped samples by site of origin. Within these groups we calculated the frequency of predicted peak-gene connections with the frequency converted to a z-score. Subgroup-specific element-gene edges are those with a z-score >= 2 and present in >50% of all samples in any site of origin. Overall, we identified on average 535 (ranging from 94 to 1,503) specific peak-gene connections for each site of origin with the peak-gene connections that target known cancer genes prioritized for experimental validation. We further overlapped the identified cancer type specific connections with COSMIC cancer genes (47). Of the 723 cancer genes investigated, we find 214 with cancer type or subtype-specific enhancer-gene connections, among which 169 connections are novel enhancer-gene connections including both canonical and non-canonical connections (142 and 27 connections respectively) (Figure 4a). Among them, we found interesting candidates. For example, we find CYP2C8, cytochrome P450 family 2 subfamily C member 8, a protein coding gene that is highly expressed in liver hepatocellular carcinoma (LIHC) and a prognostic marker in liver and pancreatic cancer (48, 49). We identified two open chromatin peaks that regulate the CYP2C8 gene and are specific to LIHC, one is a promoter peak (TCGA_331538), the other one is an enhancer peak (TCGA_331533). We observed that while the enhancer peak can be open and connecting to CYP2C8 in patients from other cancer types (e.g., CHOL, BLCA, LUAD and THCA), these two connections are present together almost only in LIHC patients (Figure 4a). Additionally, when both the promoter and enhancer connections are predicted as present, CYP2C8 is highly expressed (Figure 4b and 4c).

### Validation of ESR1 enhancers using CRISPRi

In addition to the above cancer-type specific example, we also predicted two peaks connected to ESR1 that are cancer subtype-specific. These peaks linked to ESR1 also show different behavior within breast cancer samples. Both peaks are predicted to regulate ESR1 in ER+ breast cancer subtypes but not in the majority of HER2 or in triple negative subtypes (Figure 5a, 5b and 5c). Although ESR1 has been known as important in ER+ breast cancer patients, only 1% and 7% of ER+ cases were previously explained by copy number alterations (50-53) and somatic mutations (54), respectively. In contrast, the two predicted potential enhancer peaks together can explain the high expression of ESR1 in 93% (40/43) of ER+ samples. One of the peaks, located upstream of ESR1, we denote as “enhDistal” (TCGA_219909), while the other peak located in the intronic region of ESR1, we name “enhIntron” (TCGA_219977) (Figure 5b and 5c). The peak enhDistal is only open in ER+ samples hence it is a canonical enhancer while the peak enhIntron is open in both ER+ and ER-samples, but only predicted to regulate ESR1 in ER+ samples, thus it is considered a non-canonical enhancer. In addition, both these peaks are annotated as potential enhancers by ENCODE in the MCF-7 (ER+) cell line using H3K27ac ChIP-seq and DNase-seq signals combined (Figure 5b and 5c) (2). But these enhancer-gene links were not previously identified in this patient cohort using other approaches (1).

**Figure 5.**
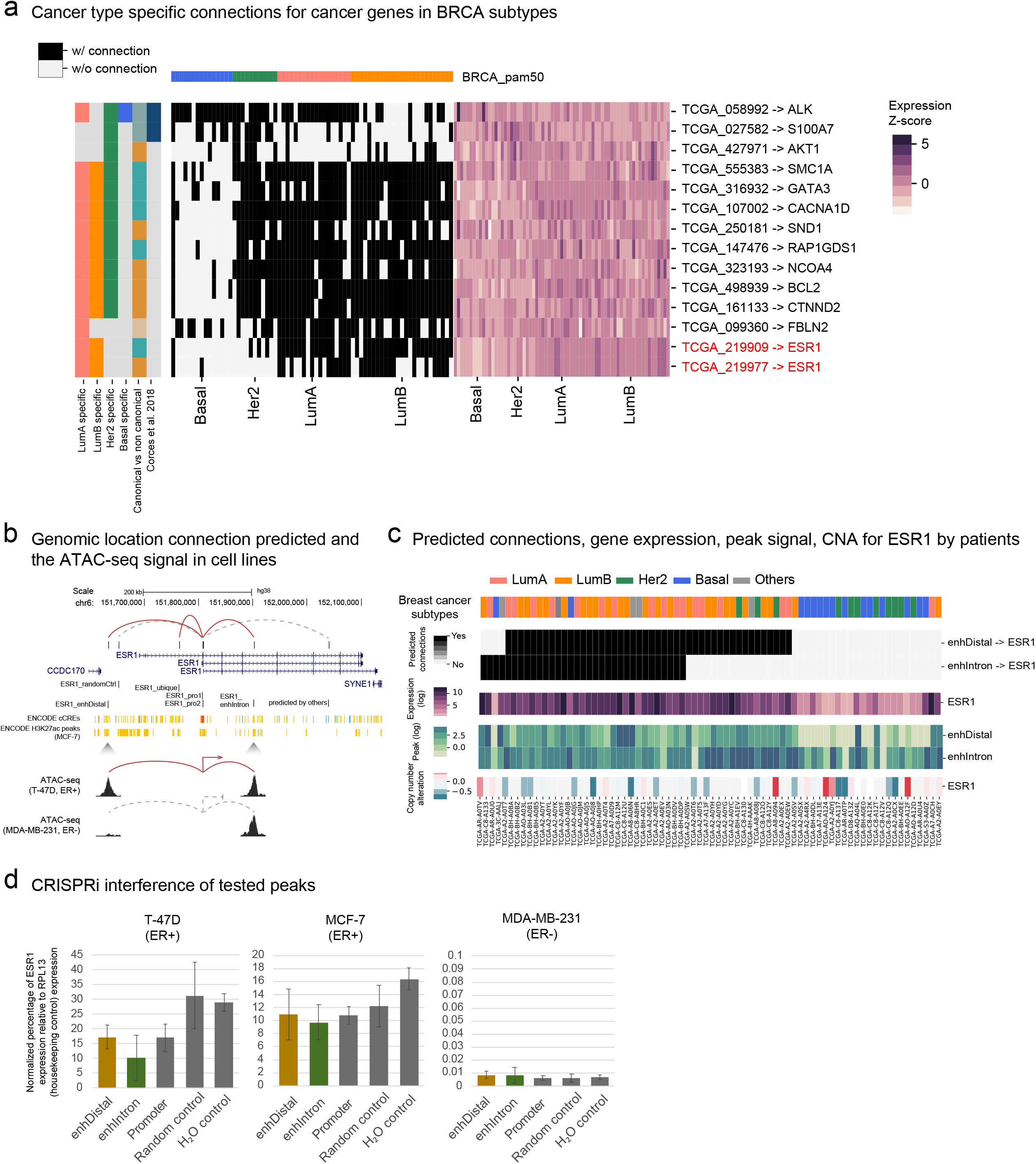
Validation of ESR1 enhancers in ER+ breast cancer. a) BRCA subtype-specific connections by subtypes for cancer genes. The left panel is the connection for each patient sample by BRCA subtypes; the right panel is the corresponding expression of genes (log, normalized by medium of the row) involved in the connection for each patient by cancer types. Vertical color bar annotates the subtype specificity for each subtype, canonical vs. non-canonical and if the connection was identified from Corces et al. 2018. The horizontal color bar annotates the subtype information for each patient. b) Genomic location of the rested enhancers, promoter control and random peak control relative to the ESR1. c) presence of enhDistal (TCGA_219909) and enhIntron (TCGA_219977) in BRCA patient cohorts, column colors represent each subtype. In addition, the expression and copy number alteration of ESR1 and peak signal by each patient. d) CRISPRi interference of tested peaks, and the normalized percentage of ESR1 expression relative to housekeeping control RPL13 expression in ER+ and ER-cell lines.

To experimentally validate these enhancers, we first generated ATAC-seq and RNA-seq data for 8 breast cancer cell lines spanning the four major breast cancer subtypes, Luminal A (LumA), Luminal B (LumB), Her2-enriched (Her2+) and Basal/Triple-negative like (TNBC). (Supplementary table 5), and applied our model on those cell lines to predict the peak-gene connections for each of them (**Methods**). ATAC-seq confirmed that both enhIntron and enhDistal are open in ER+ cell lines while enhDistal is closed in ER-cell lines. Consistent with the patient data, our model predicted that both enhancers connect to ESR1 only in ER+ cell lines, not ER-cell lines (Supplementary Figure 5).

We then tested the enhancer function of these two predicted enhancers using CRISPRi (55) in T-47D (ER+) and MDA-MB-231 (ER-) cell lines. If these enhancers function as predicted, silencing them should only decrease the expression of ESR1 in T-47D. In addition to the guides targeting enhDistal and enhIntron, we also designed two negative controls and a positive promoter control. One of the negative control guides targets a peak predicted by another method (1) to regulate ESR1 but not predicted in our approach; the other one is also an open peak, and located at a similar distance from ESR1 but not predicted to regulate ESR1 or any other genes by our model (Figure 5b, Method). Our experiments show that in T-47D cell lines, guides targeting both enhDistal and enhIntron produce a ∼2-fold decrease in ESR1 expression in MCF7 and T47D cells relative to a water control, with comparable or lower expression than guides targeted at the promoter (Figure 5d). Importantly, guides targeted at the equidistant peak have no effect on ESR1 expression while the peak predicted by the other method had no effect on ESR1 expression (Figure 5d). These results validated the function of enhDistal and enhIntron as two enhancers active in ER+ cell lines, but not ER-cell lines despite the enhIntron peak being open in ER-cell lines. In both peaks, we find no effect on ESR1 expression or cell growth in MDA-MB-231 but significant reductions in growth rate for both T47D and MCF7 in keeping with the importance of ESR1 in ER+ breast cancer.

## Discussion

Our approach, DGTAC, has demonstrated that a model trained on 3D chromatin conformation data is able to recapitulate the connections between active regulatory elements and target genes using only ATAC-seq and RNA-seq as input. It identifies non-canonical connections, which differentiate active vs. poised enhancers in a patient-specific manner. DGTAC successfully identified novel enhancers regulating important cancer genes, ESR1, which explains the high expression of ESR1 in 93% of ER+ breast cancer subtypes and was validated using CRISPRi.

Identifying enhancer-gene connections has remained a challenge in the field and it is specifically difficult to do in the absence of H3K27ac ChIP-seq or HiChIP as input. These assays are not feasible on a large scale for patient biopsies. DGTAC, however, requires only the ATAC-seq and RNA-seq to predict the enhancer-gene connection for given new samples. This enables its application to both primary and metastatic cancer patient samples when biopsies available for assays are limited. Another major advantage of our method relative to simple correlation-based approaches is that it is not limited to single element-gene correlations, instead, we utilize the combinatorial effect of multiple regulatory elements to the target gene (17). This way, we are not excluding connections that are not strongly correlated globally across all the samples for further discovery. In addition, the residual term provides the patient specific information on the effect of enhancers to the gene expression from each specific patient sample. Therefore, we can detect connections in a sample-specific manner. Currently, DGTAC was developed on the peak set from 371 primary cancer samples from TCGA (*1, 56*) and in this data set some cancer types contain less than 5 patient samples, e.g. 2 samples from CHOL and CESC, 4 samples from SKCM. The small sample size of these cancer types may not be able to capture all the representative open chromatin regions. However, since the 3D conformation of chromatin learned by DGTAC is independent of cancer type, DGTAC can still be applied to new samples where new peaks are introduced.

The identified patient-specific enhancer-gene connections provide the enhancer landscape to study the non-coding genetic variants at enhancers. In addition, the identified enhancer-gene connections make it possible for constructing more precise transcription factors (TFs) to gene networks for the 371 primary cancer patients to facilitate the discovery of important TFs and novel tumorigenesis mechanisms. DGTAC is available online at https://github.com/khuranalab/DGTAC and a docker image has been made available for download.

## Methods

### Data processing

ATAC-seq of 400 primary cancer patients was from Corces et al. (*57*) and RNA-seq data were extracted from NCI Genomic Data Commons (GDC) (*58*) with matching sample submitter_id. The ATAC-seq counts and RNA-seq counts used in this study were pre-processed by the above studies, Log2 of Transcripts Per Million (TPM) was adopted for RNA-seq counts, while log2 counts are used for ATAC-seq counts. Copy number segments of patients were obtained from GDC with matching sample submitter_id as well.

### DGTAC

ElasticNet regression was performed for all peaks within the +/-0.5 Mbp distance of TSS of each gene. The coefficients were collected as peaks’ weight to the target gene expression. For each peak of each gene, we calculate the residual (e) term representing the contribution of the single peak to the target gene expression. Peaks with zero coefficient were filtered out by giving them an artificial large residual e=10. Besides inspecting the peak itself as a single unit, we partitioned each 500 bp peak into five 100 bp windows and repeated the same regression for each of them independently and output the coefficients to calculate the residual for each of those windows. This is to consider the potential effect stemming the presence of TF footprints in individual peaks which is likely indicative of active enhancers (*59-61*). Together with peak height, window heights, peak residual and window residuals, we constructed a feature matrix for each patient including the above peak features as well as gene expression, distance to the TSS and copy number alteration of the gene for each peak-gene pair. At last, we utilized ChIA-PET of CTCF/cohesin defined loops of an MCF-7 cell line (22) to identify true peak-gene interacting pairs in a matching ER+/HER2-lumA breast cancer patient. The negative peak-gene interacting pairs were chosen by pairs that are outside of the loop. The average positive: negative ratio is 1.58:1.

By using that patient matrix, we trained a random forest model using the above set (in total 26,989 peak-gene pairs) and applied it to all 371 primary cancer patients. The model achieved an AUROC of 0.91 and AUPRC 0.92. In total, we predicted 61,502 peak-gene interacting links from 371 primary cancer patients. By evaluation of the features importance of the model, besides the “distance to TSS’’ which ranks first as the most important feature, the residual term ranks as the second important feature, ahead of gene expression, copy number alteration and peak signal itself. The same trend persists for each 100 bp peak window too (Supplementary figure 1dc), further showing the additional information provided by the residuals.

### Apply DGTAC on cell lines from ENCODE

We download the ATAC-seq bam files (from Michael Snyder, Stanford) and RNA-seq tpm (from Barbara Wold, Caltech) from ENCODE, and process them the same as Corces et al. (1) (details of datasets used are in Supplementary table 3). For ATAC-seq, we shifted the reads fragment end by “+” stranded +4 bp, “-” stranded -5 bp for the Tn5 offset, and then we counted the insertion sites on the peak set from 371 TCGA primary cancer patients (1), log2cpm was calculated using “cpm” (cpm(count, log=TRUE, prior.count = 5)) from edgeR (*62*) and normalized using “normalize.quantiles” from preprocessCore (*63*). The copy number alterations for cell lines were downloaded from The Cancer Cell Line Encyclopedia (CCLE) (*64*).

### ChIP-seq and ENCODE data process and analysis

To evaluate the potential activity of peaks with predicted target genes, we separate peaks into promoter and enhancer peaks. The promoter peaks are defined as in -2,500/+100bp of TSSs for all transcripts of its target gene; the enhancer peaks are the peaks excluding the promoter peaks from the all-peak set. Cell lines are matched by their closest cancer types and subtypes, and all H3K27ac ChIP-seq data (HepG2: ENCFF764VYK; HCT-116: ENCFF126HQH; HeLa-S3: ENCFF194XTD; PC-3: ENCFF046HYI; MCF-7: ENCFF008LRL) analyzed is “fold change over control” doswnload from encodeproject.org (Consortium et al. 2020), and analyzed using deepTools (Ramirez et al. 2016).

The newest ENCODE annotation for MCF-7 and HCT-116 regulatory elements were downloaded from https://screen.encodeproject.org/ (Consortium et al. 2020). The annotated elements were intersected with ATAC-seq peaks by 1 bp. To generate a comprehensive annotation for each peak, a peak with any of the element annotation for CTCF/PLS/pELS/dELS was annotated once for each of them. ELS is for peaks annotated by either pELS or dELS; H3Kme3 is for peaks annotated as both PLS and DNase-H3K4me3; DNase is for peaks annotated as any of the CTCF/PLS/pELS/dELS/DNase-only/DNase-H3K4me3; “Any elements” is for peaks with any of the above annotation except for DNase-only.

### Activity-by-contact score calculation

The Hi-C data was downloaded from GSE130916 (Achinger-Kawecka et al. 2020) and has been pre-processed with HiC-Pro (Servant et al. 2015). The data was further normalized by Knight-Ruiz (KR) matrix balancing using gcMapExplorer (Kumar et al. 2017). The ATAC-seq and H3K27ac ChIP-seq were processed using deepTools (Ramirez et al. 2016) by flanking +/- 1,250 bp from each peak. The ABC scores were calculated using all elements within 5 Mb of each gene TSS, and were plotted in log10 scale as described in Fulco et al. (Fulco et al. 2019).

### Gene ontology analysis

Gene ontology analysis was done using GOrilla (37).

### Cell Culture

AU-565 (CRL-2351), BT-474 (HTB-20), HEK 293T (CRL-11268), Hs 578T (HTB-126), MCF7 (HTB-22), MDA-MB-231 (HTB-26), MDA-MB-361 (HTB-27), SK-BR-3 (HTB-30) and T-47D (HTB-133) cell lines were obtained from ATCC. BT-474, HEK 293T, Hs 578T, MCF7, MDA-MB-231, and MDA-MB-361 were cultured in Dulbecco’s Modified Eagle Medium (DMEM); T-47D and AU-565 were cultured in RPMI Medium 1640; SK-BR-3 was cultured in McCoy’s 5A Medium. All media were obtained from Gibco and were supplemented with 10% Fetal Bovine Serum (Omega Scientific), 1% Sodium Pyruvate (Gibco), 1% Pen Strep (Gibco), 1% HEPES Buffer (Corning). T-47D media was additionally supplemented with 0.2 units/mL Bovine Insulin (Sigma Aldrich, IO516). Cells were regularly passaged and tested for presence of mycoplasma contamination (Mycoplasma PCR Detection Kit, Applied Biological Materials).

### RNA-seq and ATAC-seq data processing

RNA-seq and ATAC-seq were generated for 8 cell lines (MCF-7, T-47D, ZR-75-1, MDA-MB-361, BT-474, SK-BR-3, AU-565 and MDA-MB-231). Cells were grown to near confluence, pelleted and frozen at -80°C.

RNA was isolated using the RNeasy Plus Mini kit (74134, Qiagen) and sent to the Weill Cornell Medicine Genomics Core facility, where an Illumina TruSeq stranded mRNA Library prep was made using HiSeq 2500 and sequenced on the Illumina NovaSeq 6000 S1 Flow Cell, PE100 cycles. RNA-seq data were aligned to hg38 using STAR (Dobin et al. 2013), counts was calculated using HTSeq (Anders, Pyl and Huber 2015) and TPM was calculated as described by Wagner et al. (Wagner, Kin and Lynch 2012).

For ATACseq, nuclei were isolated from >95% viable cells in log phase growth and processed at the Weill Cornell Medicine Epigenomics Core using the OMIN-ATAC protocol as detailed in (Corces et al. 2017), and sequenced on Illumina NovaSeq 6000, SP Flow Cell PE50 cycles. ATAC-seq data was mapped to hg38 and processed using the ENCODE ATAC-seq pipeline (https://www.encodeproject.org/atac-seq/).

### CRISPRi sgRNA guide design and cloning

We used a modification of the GeCKO Target Guide Sequence Cloning Protocol to generate our clones (Sanjana, Shalem and Zhang 2014, Shalem et al. 2014). We utilized the software tool ChopChop to design guide RNA sequences for regions corresponding to 6 separate regulatory elements (Labun et al. 2019). 5 elements were enhancer-like elements 500 bp in size and the remaining element consisted of the region 2,500 bp upstream and 100 bp downstream of the TSS of ESR1. Exact locations and sequences are provided in Supplementary table 2. In brief, the sgRNA guide oligos targeting our regions of interest were selected to span the fullest width of the regulatory element possible while also maintaining a high guide efficiency score. For each peak, we found a 20 base pair sequence upstream of an NGG PAM sequence, which was identified as the “Forward” sequence. Each forward sequence and its reverse complement were synthesized with CACCG at the 5 ‘end of the forward strand and AAAC at the 5 ‘end of the reverse complement strand in order to make them compatible for cloning using BsmbI. The complete list of oligos used to make the guide clones is found in Table 1. All oligos were ordered using standard desalting conditions. The pCC_09 (139094, Addgene) vector (Legut et al. 2020) was digested with BsmBI (FD0454, Thermo Fisher) and treated with alkaline phosphatase (EF0651, Thermo Fisher) at 37°C for 30 min., gel-purified on a 2% agarose gel, and the digested plasmid was extracted using the QIAquick Gel Extraction Kit (28704, Qiagen). Oligo pairs were phosphorylated, annealed, and ligated with 50ng BsmBI digested plasmid. Ligated plasmids, at least 3 guide clones for each region of interest, were transformed into Stbl3 bacteria (C737303, Invitrogen), grown on Ampicillin selection and plasmid DNA was extraction using HiSpeed Plasmid maxi Kits (12662, Qiagen). Plasmid DNA was quantitated using NanoDrop spectrophotometer.

### Production of Lentivirus

Lentivirus was produced by Lipofectamine 3000 (L3000-015, Invitrogen) transfection of HEK293T cells with pCC_09 individual sgRNA clones, packaging plasmid psPAX2 (12260, Addgene) and envelope plasmid pMD2.G (12259, Addgene). Media was changed 24 hours post-transfection and virus was harvested 72 hours post-transfection. Individual viral supernatants were spun, aliquoted and frozen at -80°C until transduction.

### Lentivirus transduction

Cells to be transduced with CRISPRi guide clones were harvested in log phase and plated at 500×105 cells per well in 6 well plates with growth media plus 10ug/ml Polybrene (SCBT, sc134220). Lentivirus, at 500ul/well was added and 18 hours post-transduction, cells were split 1:2 and Puromycin was added for selection against non-transduced cells. Puromycin concentrations were determined by titration for each cell line, and ranged between 0.5-2 ug/ml. To correct for the transduction bias in MCF7 and T-47D (Hines, Yaswen and Bissell 2015) the following changes were made to the protocol. For MCF7, 1ml of virus was used to transduce 500×105 cells, with the virus was removed no sooner than 28 hours post transduction, and Puromycin was added the following morning. For T-47D, the virus concentration was not increased from the standard 500ul/well, but the virus was removed no sooner than 28 hours post-transduction and Puromycin was added 1-2 hours after the cells were re-plated. For each transduction, one well of non-transduced cells were subjected to the selection antibiotic to confirm transduction. Cells were maintained on selection antibiotic until they were frozen and/or pelleted for further analysis.

### Quantitative real-time PCR

MCF7, T-47D, SK-BR-3, MDA-MB-231 and BT-474 were transduced with the CRISPRi guide clones described above. RNA was isolated from frozen cell pellets using the Qiagen RNeasy plus Mini Kit (74134, Qiagen) and quantitated using NanoDrop. cDNA was synthesized from 1ug RNA using the qScript cDNA SuperMix (101414-106, Quantabio) according to product protocol. Transcript from 15ng cDNA was quantitated with LightCycler 480 SYBR Green I (Roche, 04887352001) using the Roche LightCycler 480 II machine. The qPCR primers used are listed in Supplementary table 2.

Expression was evaluated relative to RPL13a in individual replicates then normalized to the water/non-targeting control to evaluate ESR1 knockdown.

## Supplementary Materials

**Supplementary Figure 1**. a) accumulated counts of peak-gene connections by increasing the number of patients. b-c) clustering of BRCA subtypes and lung adenocarcinoma (LUAD),lung squamous cell carcinoma (LUSC) by principal component analysis (PCA) d) cumulated feature importance of the random forest model.

**Supplementary figure 2**. Peaks with predicted target genes are active enhancers. a) For each cancer type or subtypes, the union of all peaks with H3K27ac ChIP-seq signal for the union of all peaks with each cancer type cohorts, comparing between open peaks with predicted target gene connections vs. open peaks without predicted target gene connections vs. all other peaks closed in this cancer type cohorts. b) For both MCF-7 and HCT-116, among the open peaks with ENCODE III SCREEN annotation, the percentage of them with vs. without target gene predicted for each regulatory element category (PLS: promoter-like signatures; pELS: proximal enhancer-like signatures; dELS: distal enhancer-like signatures; ELS: enhancer-like signatures which are the union of pELS and dELS). c) ABC scores of enhancer-gene connections for K562 from Fulco et al., comparing between their predicted True and False connections; ABC scores of enhancer-gene connections from MCF-7 calculated by us comparing between our predicted True and False connections.

**Supplementary figure 3**. Features of non-canonical connections and poised enhancers

**Supplementary figure 4**. Canonical and non-canonical connection scoring matrix

**Supplementary figure 5**. DGTAC application on in house breast cancer cell lines.

**Supplementary table 1**. Model evaluation of BRCA-model and CESC-model.

**Supplementary table 2**. Cell lines data source

**Supplementary table 3**. Curated representative canonical and non-canonical connections and the scoring matrix.

**Supplementary table 4**. Gene ontology enrichment analysis for genes involved in non-canonical connections. List of genes enriched in apoptosis as well as separately enriched in proteolysis and hydrolysis.

**Supplementary table 5**. Cell lines with RNA-seq and ATAC-seq; CRISPRi guide clone sequences and qPCR primers.

